# Reconstitution of antiviral Dicer activity in vitro reveals distinct contributions of RDE-4 dsRNA-binding motifs

**DOI:** 10.64898/2025.12.17.695002

**Authors:** Elaina P. Boyle, P. Joseph Aruscavage, Claudia D. Consalvo, Brenda L. Bass

## Abstract

In *C. elegans*, antiviral RNA interference (RNAi) relies on the coordinated activity of Dicer (DCR-1), the helicase DRH-1, and the double-stranded RNA (dsRNA)-binding protein, RDE-4, yet the domain-specific contributions of RDE-4 remain unclear. Here, we reconstituted the antiviral complex from independently purified DCR-1•DRH-1 and RDE-4 to define how RDE-4 stabilizes and activates the complex. Addition of recombinant RDE-4 restored ATP hydrolysis and dsRNA cleavage to levels previously observed with the pre-assembled complex, and time-course assays revealed that RDE-4 is essential for maintaining DCR-1•DRH-1 activity.

Mutational analysis of RDE-4 revealed that both dsRBM2 and dsRBM3, but not dsRBM1, are required for reconstituting ATP hydrolysis and cleavage. Disruption of the KKxAK motif in dsRBM2 drastically reduced dsRNA affinity and abolished catalytic rescue despite preserving robust binding to DCR-1•DRH-1. Mass photometry and pulldown assays revealed that RDE-4 primarily forms DCR-1 containing complexes, predominantly through interaction with dsRBM3, with no evidence for stable interaction with DRH-1 alone. Functionally, RDE-4 enhanced DRH-1-driven ATP hydrolysis on both 52 and 106 base-pair dsRNAs, but cleavage efficiency showed strong length dependence, implicating dsRNA substrate length as an effector in this system.

Our findings establish RDE-4 as an important stabilizer of the antiviral complex and reveal distinct roles for dsRBM2 and dsRBM3 in ATP hydrolysis and dsRNA cleavage. Furthermore, our results suggest that substrate length modulates RDE-4 function, not just alone, but within the antiviral complex. These insights refine our understanding of antiviral RNAi in *C. elegans* and uncover regulatory mechanisms within the antiviral complex.

## Introduction

Upon viral infection, double-stranded RNA (dsRNA) accumulates in the cytoplasm and serves as a potent trigger of innate immune defenses (Chen and Hur 2022). In vertebrates, this response is dominated by interferon signaling, whereas invertebrates such as *Caenorhabditis elegans* rely almost exclusively on RNA interference (RNAi) for antiviral defense (Schuster, Miesen, and Van Rij 2019). Central to this pathway is the RNase III enzyme Dicer, which processes long viral dsRNA into short interfering RNAs (siRNAs). These siRNAs are then loaded into Argonaute proteins to mediate sequence-specific silencing. However, Dicer is also responsible for producing microRNAs (miRNAs), whose precursors differ structurally from viral dsRNA (Wilson and Doudna 2013). To accommodate diverse dsRNA substrates, some organisms encode multiple Dicers. For example, *Drosophila melanogaster* uses Dicer-1 (dmDcr-1) for miRNA maturation and Dicer-2 (dmDcr-2) for siRNA production (Lee et al. 2004; Marques et al. 2013). In contrast, *C. elegans* and *H. sapiens* encode a single Dicer. The sole *C. elegans* Dicer (DCR-1) functions in multiple RNAi pathways, and this functional versatility depends on accessory factors that modulate DCR-1’s activity and specificity (Duchaine et al. 2006; Grishok et al. 2001; Pavelec et al. 2009; Thivierge et al. 2012).

Two accessory proteins are critical for antiviral RNAi in *C. elegans*: the dsRNA-binding protein (dsRBP) *R*NAi *De*ficient-4 (RDE-4) and *D*icer-*R*elated *H*elicase-1 (DRH-1) (Ashe et al. 2015; Guo et al. 2013; Lu et al. 2005; Sowa et al. 2020; Tabara et al. 2002; Thivierge et al. 2012). RDE-4 contains two canonical dsRNA-binding motifs (dsRBMs 1 and 2) and a third degenerate dsRBM within its C-terminal domain (CTD), a feature common among dsRBM-containing proteins (Masliah, Barraud, and Allain 2012; Parker, Eckert, and Bass 2006). Interactions between Dicers and dsRBPs have been described in multiple systems, including human Dicer and its association with TRBP (human immunodeficiency virus trans-activating response RNA-binding protein) and dmDcr-2’s interaction with R2D2 (Hansen et al. 2019; Liu et al. 2018; Yamaguchi et al. 2022). In both cases, structural studies show that the degenerate dsRBM, or CTD, of the dsRBP contacts the Hel2i subdomain of Dicer’s helicase. Similarly, cryo-EM studies of the *C. elegans* antiviral complex revealed density consistent with a single domain of RDE-4 bound to the helicase of DCR-1, though the resolution was insufficient to define the interface (Consalvo et al. 2024). By analogy, this interaction may involve the degenerate dsRBM of RDE-4 engaging with the Hel2i subdomain. However, DRH-1 also contains a helicase domain of the same family as that in Dicer (Fig. 1A), raising the possibility that RDE-4 interacts with DCR-1, DRH-1, or both. Given the inherent flexibility of dsRBMs, which often prevents their visualization by structural techniques such as cryo-EM (Jouravleva et al. 2022; Lee et al. 2023; Su et al. 2022), biochemical approaches are required to define these interactions with higher resolution.

**Figure 1.**
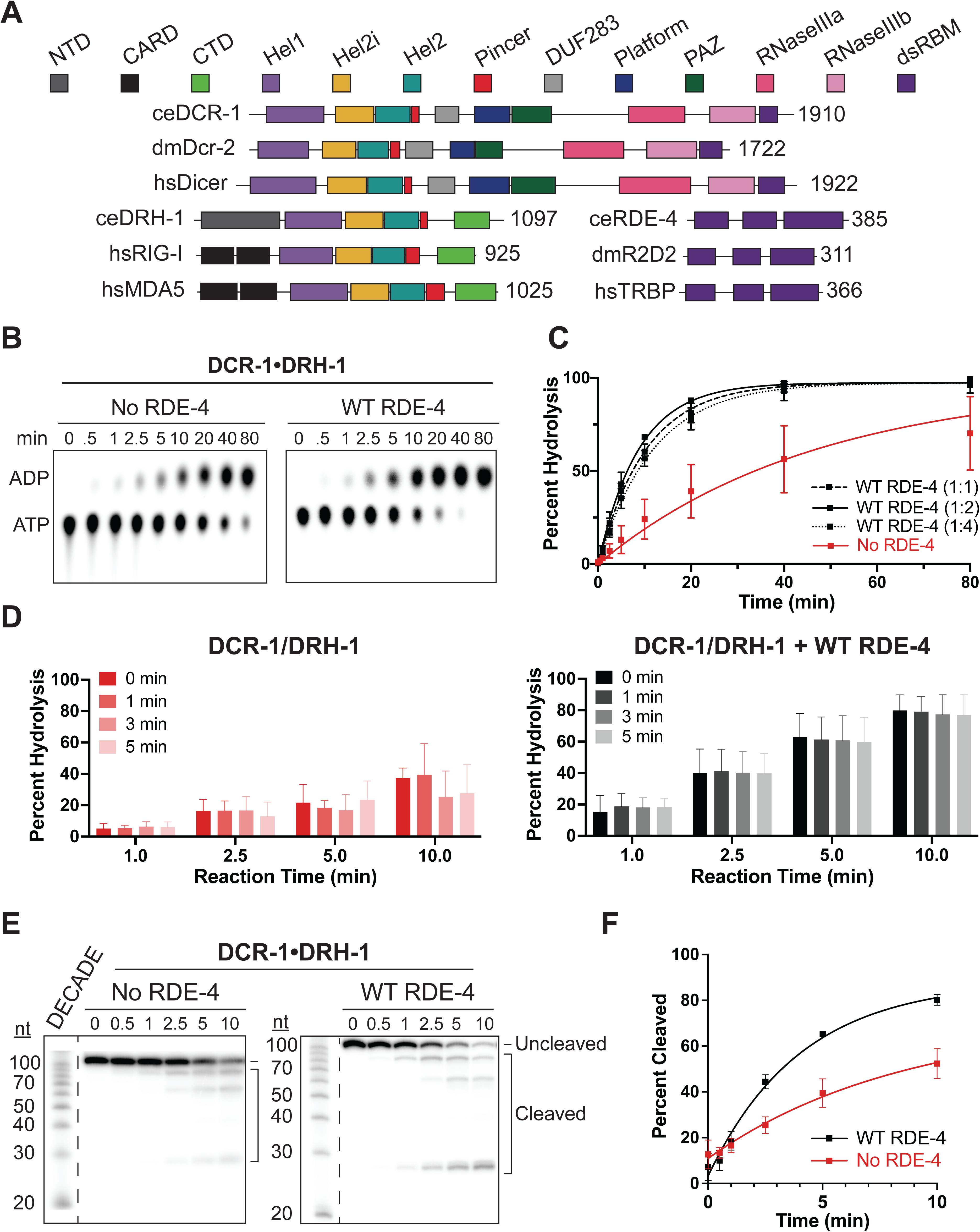
Combining DCR-1•DRH-1 with RDE-4 reconstitutes activity of the antiviral complex. A. Adapted from (Consalvo et al. 2024). Conserved domains are depicted as colored rectangles corresponding with the key above. NCBI conserved domains, AlphaFold, and previous reports were used in defining domain boundaries (Consalvo et al. 2024; Hansen et al. 2019; Jumper et al. 2021; Lu et al. 2020; Parker et al. 2008; Varadi et al. 2024). Numbers to the right of the open-reading frame represent the number of amino acids in the protein. Species: ce, *C. elegans*; dm, *Drosophila melanogaster* and hs, *Homo sapiens*. B. DCR-1•DRH-1 (150 nM) and RDE-4 (300 nM) were incubated with 106 BLT dsRNA (400 nM) and α-^32^P-ATP at 20°C. Percent ATP hydrolysis was monitored using thin-layer chromatography (TLC). A representative PhosphorImage is shown, and positions of ATP and ADP are indicated. C. Quantification of multiple ATP hydrolysis assays as shown in panel B. Data points are mean ± SD (n=2) with minus RDE-4 in red and plus RDE-4 in black. Ratios of DCR-1•DRH-1 to RDE-4 are indicated. D. Quantification of ATP hydrolysis assays as in C, but with pre-incubation to test heat stability of proteins. DCR-1•DRH-1 (left) or DCR-1•DRH-1 and RDE-4 (right) with 1:2 fold excess RDE-4 were preincubated for times indicated in inset, prior to addition of 106 BLT dsRNA and α-^32^P-ATP and incubation for reaction times as indicated on x-axis. All incubations were at 20°C. Percent ATP hydrolysis was monitored using TLC. Data points are mean ± SD (n=3). See also Materials and Methods and Supplemental Figs. S2A-C. E. Single-turnover cleavage assays of DCR-1•DRH-1 (10 units) and RDE-4 (2-fold molar excess) with 106 BLT dsRNA (1 nM) and ATP (5 mM) at 20°C for times indicated (see Materials and Methods). Sense (top) strand was 5’ ^32^P-end labeled. Products were separated by 17% denaturing PAGE, and a representative PhosphorImage is shown (n≥3). Decade RNA markers were loaded in far left lane of each gel, with nucleotide (nt) length to left of gel. Empty lane was removed for clarity; boundaries of gel-splicing indicated by dashed line. F. Quantification of multiple single-turnover assays as shown in panel E, with labels to the right of each curve. Data points are mean ± SD (n=3).

Previous studies show that RDE-4 enhances dsRNA binding and promotes both ATP hydrolysis and cleavage by the DCR-1•DRH-1 complex (Consalvo et al. 2024). Genetic and biochemical experiments suggest that dsRBM2 and the CTD (containing dsRBM3) are particularly important, but these studies were conducted either in cell extracts or in vivo, where additional factors may influence complex assembly and function (Blanchard et al. 2011; Parker et al. 2006, 2006). Thus, the precise contributions of individual RDE-4 domains, and whether they act primarily through RNA binding or protein-protein contacts, remain unclear.

Here, we reconstitute the antiviral complex using recombinant proteins purified from insect cells and systematically dissect the roles of RDE-4 domains in stabilizing and activating DCR-1•DRH-1. By combining activity assays, dsRNA binding measurements, pulldowns, and mass photometry, we define how RDE-4 promotes antiviral activity and identify the distinct contributions of dsRBM2 and dsRBM3 to complex function.

## Results

### Combining DCR-1•DRH-1 with RDE-4 reconstitutes activity of the antiviral complex

To investigate the role of RDE-4 in the DCR-1•DRH-1•RDE-4 antiviral complex, we expressed DCR-1•DRH-1 and RDE-4 separately in Sf9 cells using baculovirus vectors. Recombinant DCR-1•DRH-1 was purified to ∼81% homogeneity (Supplemental Fig. S1A). To maintain consistency between recombinant proteins, RDE-4 constructs previously expressed in *S. cerevisiae* (Parker et al. 2006; Parker, Maity, and Bass 2008) were instead produced using the baculovirus expression system. Wild-type RDE-4 was purified to ∼72% homogeneity, and its sequence was verified by mass spectrometry (Supplemental Figs. S1A-B).

Previous work characterized the pre-assembled antiviral complex, establishing parameters for ATP hydrolysis, dsRNA binding, and cleavage, using a 106 base pair (bp) dsRNA substrate (Consalvo et al. 2024). In these assays, the antiviral complex was purified after co-expression of all three proteins, and thus, pre-assembled before adding other reaction components. To compare activities to a complex reconstituted by separately adding RDE-4 to DCR-1•DRH-1, we first performed ATP hydrolysis assays (Figs. 1B-C). An equimolar amount of DCR-1•DRH-1 and RDE-4 were combined, and to ensure complete saturation of binding, RDE-4 was also tested at two- or four-fold molar excess of DCR-1•DRH-1, though analytical ultracentrifugation revealed a 1:1:1 ratio of DCR-1, DRH-1 and RDE-4 in the pre-assembled complex (Consalvo et al. 2024). We were able to achieve complete reconstitution of ATP hydrolysis with a plateau around 90%. Additionally, k_obs_ values correlated well with those previously obtained for the pre-assembled complex (Table 1). However, throughout data collection there were indications that the DCR-1•DRH-1 complex was unstable without RDE-4. For example, the DCR-1•DRH-1 complex was far less amenable to purification than the DCR-1•DRH-1•RDE-4 complex. Additionally, if the protein was left on ice before beginning an assay, a noticeable decrease in activity was observed.

**Table 1.** Reconstitution of cleavage and hydrolysis.

| | $K_{obs}$ ($\text{min}^{-1}$)<br>(ATP Hydrolysis) | $v_0$ (%/min)<br>(ATP Hydrolysis) | $K_{obs}$ ($\text{min}^{-1}$)<br>(Cleavage) | $v_0$ (%/min)<br>(Cleavage) |
| --- | --- | --- | --- | --- |
| DCR-1•DRH-1•RDE-4* | $0.11 \pm 0.01$ | -- | $0.14 \pm 0.01$ | -- |
| DCR-1•DRH-1 + RDE-4 (1:1) | $0.096 \pm 0.005$ | $8.7 \pm 0.3$ | -- | -- |
| DCR-1•DRH-1 + RDE-4 (1:2) | $0.12 \pm 0.006$ | $8.8 \pm 0.6$ | $0.25 \pm 0.03$ | $16 \pm 1$ |
| DCR-1•DRH-1 + RDE-4 (1:4) | $0.088 \pm 0.006$ | $6.8 \pm 0.5$ | -- | -- |

\*(Consalvo et al. 2024)
-- not determined.

To address concerns that plateaus for DCR-1•DRH-1 reactions were indicative of the complex losing activity over time, rather than saturation of the enzyme, DCR-1•DRH-1 was pre-incubated at the reaction temperature for various times before adding a master mix containing reaction buffer, ATP and dsRNA (Fig. 1D and Supplemental Fig. S2A-C). These experiments revealed that DCR-1•DRH-1 was indeed losing activity over time. For example, the percent hydrolysis measured at 10 minutes decreased in samples pre-incubated for over 1 minute (Fig. 1D, left). In contrast, addition of RDE-4 stabilized the complex, demonstrated by uniform hydrolysis over the 10-minute assay, independent of pre-incubation time (Fig. 1D, right). After testing various conditions, we concluded that DCR-1•DRH-1 was stable without RDE-4 for ∼10 minutes at reaction temperature. Accordingly, all subsequent time courses were limited to 10 minutes to enable reliable comparisons between reactions with and without RDE-4. Additionally, units of activity for DCR-1•DRH-1 were established to maintain consistency in activity levels, independent of purification batch (see Materials and Methods).

Activity of the reconstituted antiviral complex was further investigated using cleavage assays with 106-bp ^32^P-radiolabeled dsRNA (Figs. 1E-F). Cleavage products were visualized after denaturing polyacrylamide gel electrophoresis (PAGE), and the fraction of cleaved dsRNA relative to total radioactive dsRNA signal was quantified. Conditions with two-fold excess RDE-4 relative to DCR-1•DRH-1 were chosen for subsequent experiments to favor DCR-1•DRH-1•RDE-4 complex formation. Previous reports led to concerns about excess RDE-4 leading to cleavage inhibition (Parker et al. 2006), however this was not observed with 2-fold excess. Instead we observed a robust improvement in cleavage upon addition of RDE-4 (Figs. 1E-F). Trends agreed well with the previous characterization of the antiviral complex where DCR-1•DRH-1+RDE-4 plateaus around 90% cleavage, and DCR-1•DRH-1 alone plateaus around 60% cleavage (Figs. 1E-F) (Consalvo et al. 2024). Differences in cleavage pattern can be attributed to the change in overall protein concentration, and the lack of a cyclic phosphate on the dsRNA which allows cleavage from either end (See Materials and Methods).

### dsRBM2 and dsRBM3 are essential for catalytic reconstitution, whereas dsRBM1 is dispensable

After establishing conditions under which WT RDE-4 restored ATP hydrolysis and cleavage activity, we designed and expressed mutant RDE-4 constructs to define the domains required for catalytic reconstitution. Two truncation constructs were designed to separate the effects of dsRBMs 1 and 2 from dsRBM3 (Fig. 2A). Truncation R_1_•R_2_ was purified to ∼64% homogeneity and truncation R_3_ to ∼62% homogeneity (Supplemental Figs. S1A-C). Additionally, point mutations were introduced into the KKxAK motif of dsRBMs 1 and 2, a region known to be important for dsRNA binding and conserved in dsRBMs of other proteins (Ryter 1998; St Johnston et al. 1992; Tian et al. 2004) (Figs. 2A-B). dsRBM point mutant constructs R_1_ ^AA^•R_2_ •R_3_ , R_1_ •R_2_ ^AA^•R_3_ and R_1_ ^AA^•R_2_ ^AA^•R_3_ were purified to ∼53%, 76% and 67% homogeneity, respectively, and all protein sequences were confirmed using mass spectrometry (University of Utah Mass Spectrometry Core Facility, see Supplemental Figs. S1A-B).

**Figure 2.**
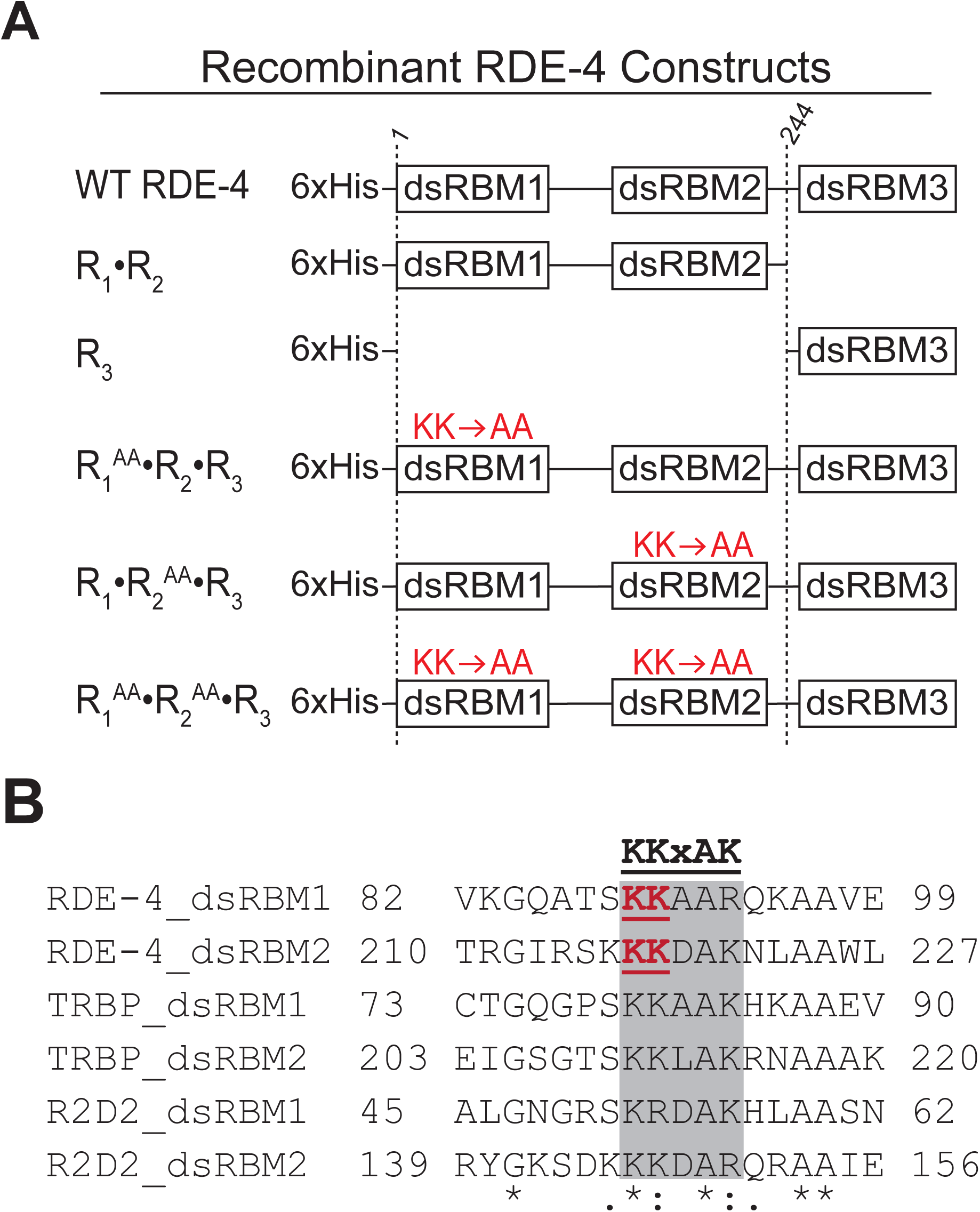
Mutational analysis of RDE-4. A. Domain structures of RDE-4 constructs. Numbers at top of dotted lines indicate amino acid positions relative to WT RDE-4 (no tag). Amino acid changes made in dsRBM are shown in red. B. Multiple sequence alignment of dsRBMs in RDE-4 and other dsRBPs, highlighting KKxAK motif in grey. The lysine to alanine mutations were K89A/K90A in dsRBM1 and K217A/K218A in dsRBM2 (shown in red). An asterisk (*) indicates positions which have a single fully conserved residue. A colon (:) indicates conservation between groups with strongly similar properties. A period (.) indicates conservation between groups with weakly similar properties (Sievers et al. 2011).

Each mutant was tested in ATP hydrolysis assays using 106-bp and 52-bp dsRNA substrates with blunt termini to mimic viral substrates (Figs. 3A-B and Supplemental Fig. S3A)(Welker et al. 2011), and kinetic parameters are summarized in Table 2. For comparison, reactions containing no RDE-4 or WT RDE-4 (2-fold molar excess) were included. Strikingly, most mutants failed to restore ATP hydrolysis to WT levels. Truncation R_1_•R_2_ failed to restore WT activity, resembling the no RDE-4 control, consistent with the proposed requirement for dsRBM3-mediated interaction with DCR-1•DRH-1 in vivo or with extracts (Blanchard et al. 2011; Parker et al. 2006, 2008). However, truncation R_3_ also failed to restore WT activity, indicating that this interaction alone was insufficient to enhance ATP hydrolysis. Mutations in the dsRBM2 KKxAK motif (R_1_•R_2_ ^AA^•R_3_ and R_1_ ^AA^•R_2_ ^AA^•R_3_) also precluded rescue by RDE-4, highlighting a key role for dsRBM2. In contrast, mutation of dsRBM1 (R_1_ ^AA^•R_2_ •R_3_) had no effect, and this construct restored ATP hydrolysis to WT levels. Differences in primary data became particularly evident after 2.5 minutes; WT RDE-4 and R_1_ ^AA^•R_2_ •R_3_ display a noticeable reduction in the ATP spot together with a darker ADP spot, while all other mutants resembled the no RDE-4 control (Figs. 3A-B). Thus, dsRBM1 is dispensable, whereas dsRBM2 and dsRBM3 are required for reconstitution of catalytic activity. Results were consistent across assays using both 106-bp and 52-bp dsRNA substrates (Table 2, Figs. 3A-B and Supplemental Fig. S3A).

**Figure 3.**
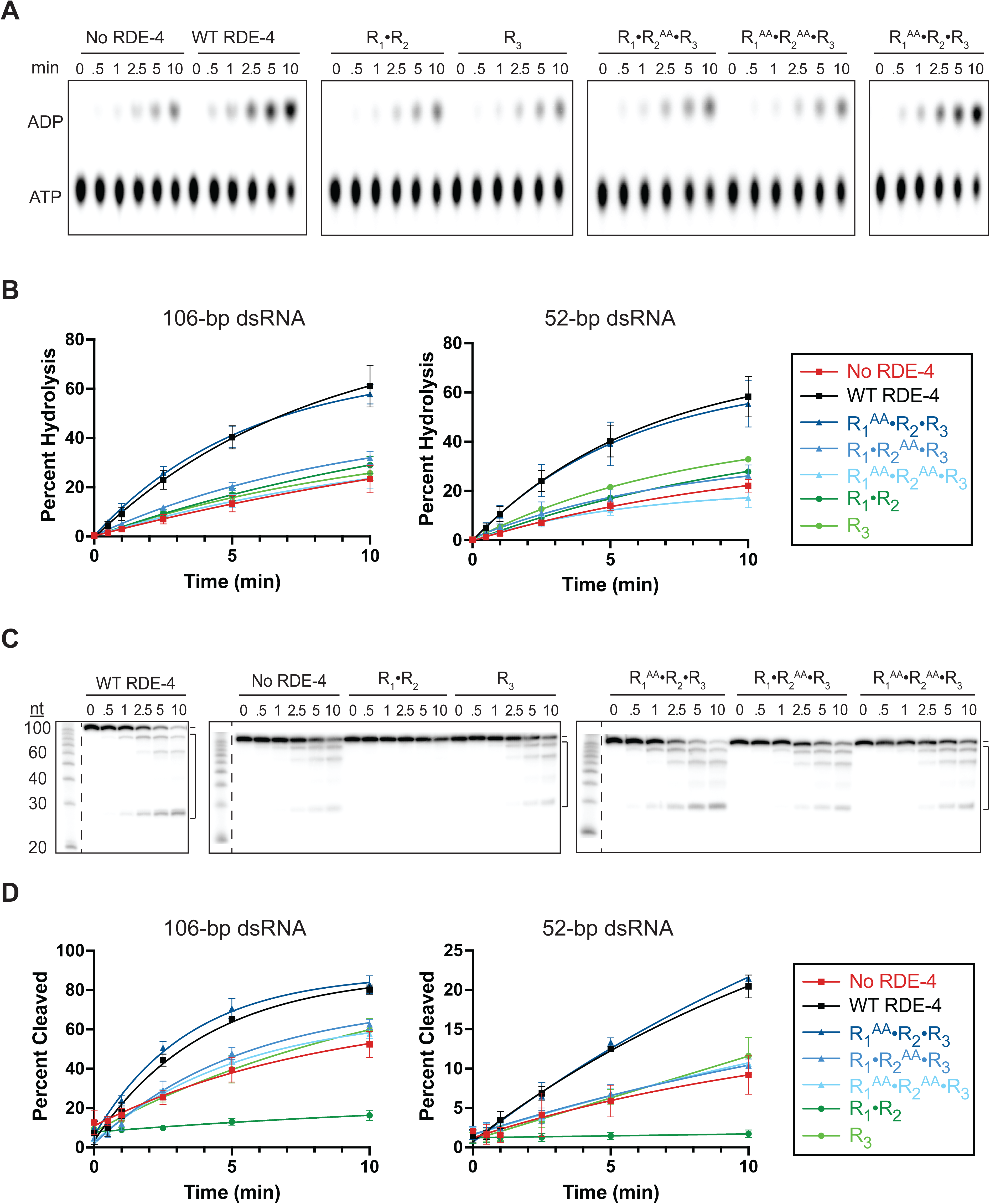
dsRBM2 and dsRBM3 are essential for catalytic reconstitution, whereas dsRBM1 is dispensable. A. DCR-1•DRH-1 (10 units) and RDE-4 or mutants (2-fold molar excess) were incubated with 106 BLT or 52 BLT dsRNA (400 nM) and α-^32^P-ATP at 20°C. Percent ATP hydrolysis at times indicated was monitored using TLC. Representative PhosphorImages with 106 BLT are shown with positions of ATP and ADP mobility indicated (n≥3). See also Supplemental Fig. S3A for primary data with 52 BLT dsRNA. B. Quantification of multiple ATP hydrolysis assays as in panel A. Data points are mean ± SD (n=3) with legend on the right. C. Single-turnover cleavage assays of DCR-1•DRH-1 (10 units) and RDE-4 or mutants (2-fold molar excess), with 106 BLT or 52 BLT dsRNA (1 nM) and ATP (5 mM) at 20°C. Sense strand was 5’ ^32^P-end labeled. Products were separated by 17% denaturing PAGE and representative PhosphorImages are shown for 106 BLT dsRNA (n=3), with size markers on left of gel. Gels were cut between lanes (as marked by dotted lines) to remove empty wells. No other image manipulations were performed. See also Supplemental Fig. S3B for primary data with 52 BLT dsRNA. D. Quantification of single-turnover assays in panel C. Data points are mean ± SD (n=3) with legend on the right. Y-axes in panel D are set to different ranges so that differences between mutants are more visible.

**Table 2.** Effects of RDE-4 mutants on cleavage and hydrolysis activity.

| | $K_{obs}$ ($\text{min}^{-1}$)<br>(Cleavage) | $K_{obs}$ ($\text{min}^{-1}$)<br>(ATP Hydrolysis) | $v_0$ (%/min)<br>(Cleavage) | $v_0$ (%/min)<br>(ATP Hydrolysis) |
| --- | --- | --- | --- | --- |
| **DCR-1•DRH-1** |  |  |  |  |
| 106 bp dsRNA | $0.12 \pm 0.06$ | $0.048 \pm 0.05$ | $9.4 \pm 0.2$ | $2.7 \pm 0.05$ |
| 52 bp dsRNA | $0.075 \pm 0.1$ | $0.093 \pm 0.03$ | $0.95 \pm 0.4$ | $2.8 \pm 0.1$ |
| **DCR-1•DRH-1 + WT RDE-4 (1:2)** |  |  |  |  |
| 106 bp dsRNA | $0.25 \pm 0.03$ | $0.13 \pm 0.03$ | $16 \pm 1$ | $9.1 \pm 0.2$ |
| 52 bp dsRNA | $0.067 \pm 0.02$ | $0.16 \pm 0.04$ | $2.3 \pm 0.3$ | $9.5 \pm 0.3$ |
| **DCR-1•DRH-1 + R<sub>1</sub>•R<sub>2</sub> (1:2)** |  |  |  |  |
| 106 bp dsRNA | $0.060 \pm 0.08$ | $0.068 \pm 0.03$ | $0.81 \pm 0.1$ | $3.6 \pm 0.06$ |
| 52 bp dsRNA | *n. r.* | $0.086 \pm 0.01$ | $-0.02 \pm 0.03$ | $3.6 \pm 0.2$ |
| **DCR-1•DRH-1 + R<sub>3</sub> (1:2)** |  |  |  |  |
| 106 bp dsRNA | $0.088 \pm 0.04$ | $0.090 \pm 0.03$ | $7.0 \pm 0.8$ | $3.4 \pm 0.01$ |
| 52 bp dsRNA | $0.00043 \pm 0.05$ | $0.13 \pm 0.009$ | $0.83 \pm 0.09$ | $4.9 \pm 0.02$ |
| **DCR-1•DRH-1 + R<sub>1</sub><sup>AA</sup>•R<sub>2</sub>•R<sub>3</sub> (1:2)** |  |  |  |  |
| 106 bp dsRNA | $0.29 \pm 0.04$ | $0.18 \pm 0.02$ | $17 \pm 1$ | $10. \pm 0.3$ |
| 52 bp dsRNA | $0.054 \pm 0.03$ | $0.18 \pm 0.05$ | $2.1 \pm 0.3$ | $9.6 \pm 0.2$ |
| **DCR-1•DRH-1 + R<sub>1</sub>•R<sub>2</sub><sup>AA</sup>•R<sub>3</sub> (1:2)** |  |  |  |  |
| 106 bp dsRNA | $0.19 \pm 0.03$ | $0.11 \pm 0.02$ | $10. \pm 1$ | $4.6 \pm 0.08$ |
| 52 bp dsRNA | $0.050 \pm 0.05$ | $0.13 \pm 0.06$ | $0.89 \pm 0.07$ | $4.2 \pm 0.3$ |
| **DCR-1•DRH-1 + R<sub>1</sub><sup>AA</sup>•R<sub>2</sub><sup>AA</sup>•R<sub>3</sub> (1:2)** |  |  |  |  |
| 106 bp dsRNA | $0.19 \pm 0.02$ | $0.079 \pm 0.04$ | $9.3 \pm 0.7$ | $2.9 \pm 0.04$ |
| 52 bp dsRNA | $0.034 \pm 0.03$ | $0.17 \pm 0.05$ | $0.86 \pm 0.07$ | $2.9 \pm 0.08$ |

n.r., not reliably determined. Italics indicate poor fit. See Materials and Methods.

We next assessed how RDE-4 mutants influenced cleavage of radiolabeled 106-bp or 52-bp dsRNA (Figs. 3C-D, Table 2 and Supplemental Fig. S3B). Because ATP hydrolysis and translocation are driven primarily by DRH-1, whereas cleavage is mediated by DCR-1 (Aderounmu et al. 2025; Consalvo et al. 2024), we anticipated that some mutants might differentially affect these processes. Instead, trends were largely consistent between the two assays. The R_3_ truncation and dsRBM2 mutants (R_1_•R_2_ ^AA^•R_3_, R_1_ ^AA^•R_2_ ^AA^•R_3_) failed to restore cleavage to WT activity and closely resembled the no RDE-4 control. In these reactions, cleavage products were present, but the uncleaved dsRNA band remained prominent at 10 minutes (Fig. 3C).

One exception was the R_1_•R_2_ truncation, which not only failed to restore cleavage but strongly inhibited it, yielding activity near background levels. This likely reflects competition for dsRNA as hydrolysis assays were performed with excess substrate, but cleavage assays were not, suggesting that R_1_•R_2_ coats dsRNA and blocks DCR-1•DRH-1 binding. Accordingly, no cleavage products were detected on gels (Fig. 3C). Finally, R_1_ ^AA^•R_2_ •R_3_ maintained cleavage at WT levels, with progressive loss of uncleaved dsRNA between 2.5 and 10 minutes, similar to the pattern observed with addition of WT RDE-4. When tested with 52-bp dsRNA, overall activity was greatly reduced (note the difference in range of y-axes in Fig. 3D). However, trends mirrored those observed with the 106-bp substrate; only R_1_ ^AA^•R_2_ •R_3_ maintained the levels observed with WT RDE-4, while R_3_ and dsRBM2 mutants resembled the no RDE-4 control, and R_1_•R_2_ inhibited cleavage completely (Table 2, Figs. 3C-D and Supplemental Fig. S3B). This difference may result from longer dsRNAs providing sufficient space to accommodate additional proteins, as described further in the Discussion.

### dsRBM point mutations alter dsRNA binding

To gain mechanistic insight into how RDE-4 mutants impacted ATP hydrolysis and dsRNA cleavage, we first conducted gel shift assays with ^32^P-radiolabeled 106-bp dsRNA (Figs. 4A-B). Values for the dissociation constant (K_d_) and Hill coefficient were determined for WT and mutant RDE-4 constructs and are summarized in Table 3.

**Figure 4.**
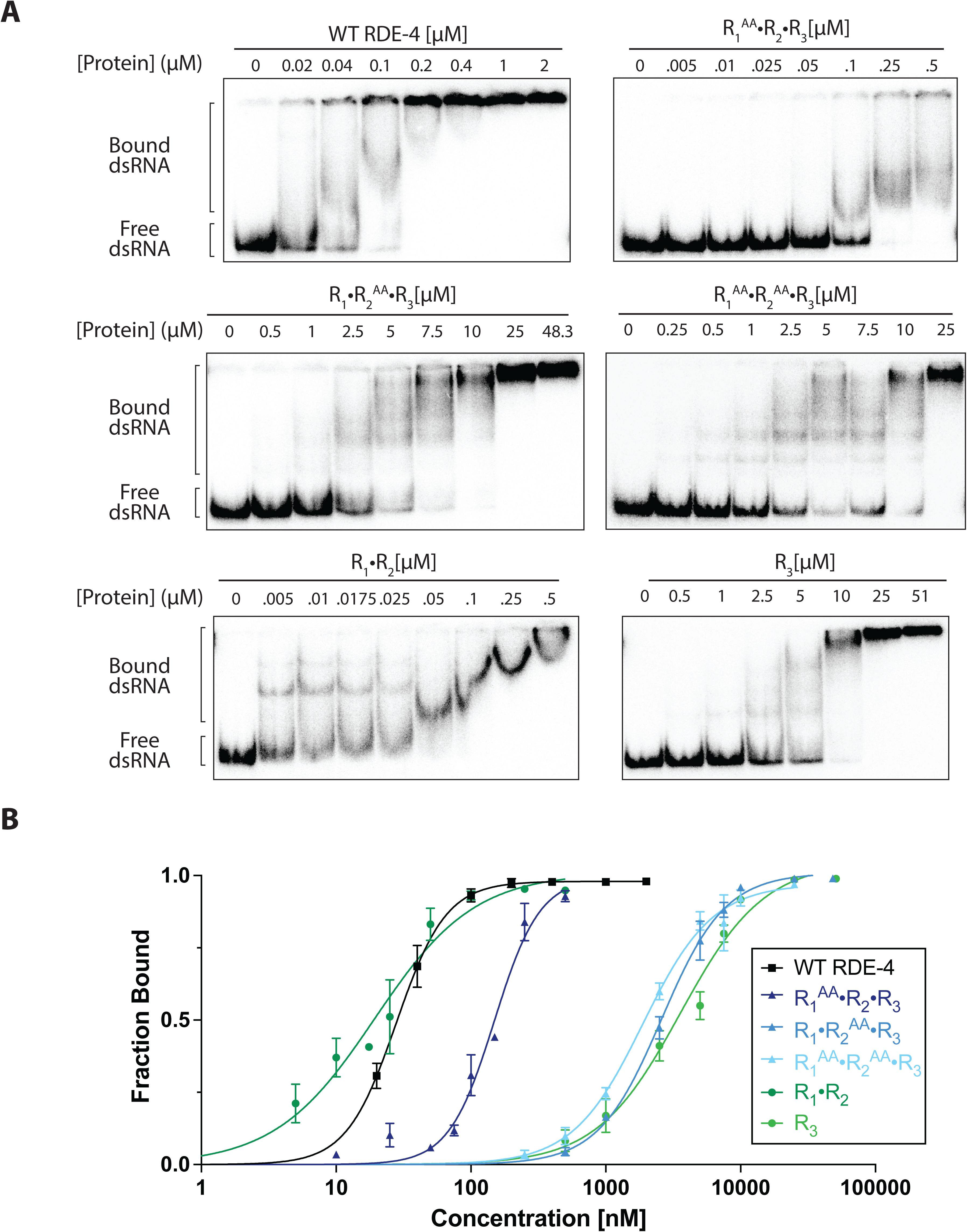
dsRBM point mutations alter dsRNA binding. A. Representative PhosphorImages showing gel mobility shift assays with increasing concentrations of RDE-4 or dsRBM point mutants, ranging from 0-51 µM with 20 pM 106 BLT dsRNA. Concentrations are total protein determined by absorbance at 280 nM and/or densitometry. Sense strand was 5’ ^32^P-end labeled. Labels on left indicate that all dsRNA that migrated through the gel more slowly than free dsRNA was considered bound. Solid boxes indicate boundaries of individual gels. B. Quantification of panel A was used to generate binding isotherms for WT and RDE-4 mutant constructs. Fraction bound = (1-[dsRNA]_free_)/[dsRNA]_total_ was fit to calculate the dissociation constant, K_d_, using Hill formalism (see Materials and Methods). Data points are mean ± SD (n=3).

**Table 3.** dsRNA binding affinity and cooperativity.

| | $K_d$ (nM) | Hill Coefficient |
| --- | --- | --- |
| WT RDE-4 | $28 \pm 0.09$ | $2.4 \pm 0.02$ |
| $R_1 \bullet R_2$ | $16 \pm 7$ | $0.84 \pm 0.4$ |
| $R_3$ | $3600 \pm 500$ | $1.4 \pm 0.2$ |
| $R_1^{AA} \bullet R_2 \bullet R_3$ | $150 \pm 20$ | $2.6 \pm 0.5$ |
| $R_1 \bullet R_2^{AA} \bullet R_3$ | $2600 \pm 80$ | $1.8 \pm 0.08$ |
| $R_1^{AA} \bullet R_2^{AA} \bullet R_3$ | $1900 \pm 100$ | $1.7 \pm 0.1$ |

The 106-bp substrate used here was not identical to the 104-bp dsRNA used in prior studies (Parker et al. 2006, 2008) but was highly comparable as both substrates have blunt termini and are perfectly base-paired dsRNAs of nearly the same length. While absolute values differ somewhat, likely due to differences in expression systems, the overall trends were consistent. For instance, WT RDE-4 and truncation construct R_1_•R_2_ showed similar affinities in both systems. The dsRBM1 point mutant R_1_ ^AA^•R_2_ •R_3_ , which was previously reported to bind with near-WT affinity, displayed reduced binding by ∼5-fold in our system. Removing both canonical dsRBMs 1 and 2 resulted in weak affinity for dsRNA, although residual binding by R_3_ was detectable at high concentrations. The dsRBM2 mutants, R_1_•R_2_^AA^•R_3_ and R_1_ ^AA^•R_2_ ^AA^•R_3_ , which have not been tested in prior studies, showed ∼90-fold and ∼70-fold reduced affinity, respectively, highlighting the critical contribution of the KKxAK motif in dsRBM2. Interestingly, simultaneous mutation of dsRBMs 1 and 2 did not exacerbate the binding defect beyond that of dsRBM2 alone, in fact, R_1_ ^AA^•R_2_ ^AA^•R_3_ bound slightly better than R_1_ •R_2_ ^AA^•R_3_ . These results align with the functional impairments observed in our ATP hydrolysis and cleavage assays.

Assessment of cooperativity by Hill coefficients revealed a similar overall pattern. WT RDE-4 and R_1_ ^AA^•R_2_ •R_3_ both bound dsRNA with high cooperativity (Hill coefficients >2). Previous work from the Bass lab indicates dsRBM2 plays a key role in promoting cooperative binding, and that constructs containing either dsRBMs 1 and 2 or dsRBMs 2 and 3 can exhibit cooperative behavior under certain conditions (Parker et al. 2008). While the precise contribution of each domain may vary depending on RNA length and context, these findings are broadly consistent with the idea that multiple dsRBMs enhance cooperative recognition. Similarly, truncation constructs R_1_•R_2_ and R_3_ showed largely non-cooperative binding (Hill coefficients ∼0.8-1.4), and the dsRBM2 mutant constructs R_1_•R_2_ ^AA^•R_3_ and R_1_ ^AA^•R_2_ ^AA^•R_3_ retained partial cooperativity (Hill coefficients ∼1.7-1.8), suggesting that while dsRBM2 is central to affinity, additional domains contribute to binding mode and cooperativity.

### dsRBM3 is the primary driver for interaction with DCR-1•DRH-1

Protein-protein interactions were first evaluated with a pulldown assay using StrepTactin-bound magnetic beads that bind the tag on DRH-1 (Fig. 5). DCR-1•(OSF)DRH-1 and WT or mutant RDE-4 were combined with 2-fold molar excess RDE-4 and incubated with StrepTactin magnetic beads (Cube Biotech). Proteins present in the input and elution were visualized using an SDS-PAGE gel stained with fluorescent dye (SYPRO-red). Percent recovery was determined from the ratio of signal of RDE-4 to DRH-1 present in the elution fraction and then normalized to WT (Figs. 5A-D). Truncation R_3_ (containing only dsRBM3) was recovered at ∼72%, whereas R_1_•R_2_ (lacking dsRBM3) was recovered at only ∼8% (Figs. 5A-B). These results support a model where dsRBM3 primarily mediates RDE-4 interaction with the antiviral complex. While dsRBM2 may contribute, these effects are likely mediated by dsRNA as cleavage data with truncation R_1_•R_2_ indicates a functional requirement for dsRBM3 to get productive interaction with the antiviral complex (Fig. 3C-D) (See Discussion). While no additional dsRNA was added to these reactions, there may be some nucleic acid contamination remaining from protein purification.

**Figure 5.**
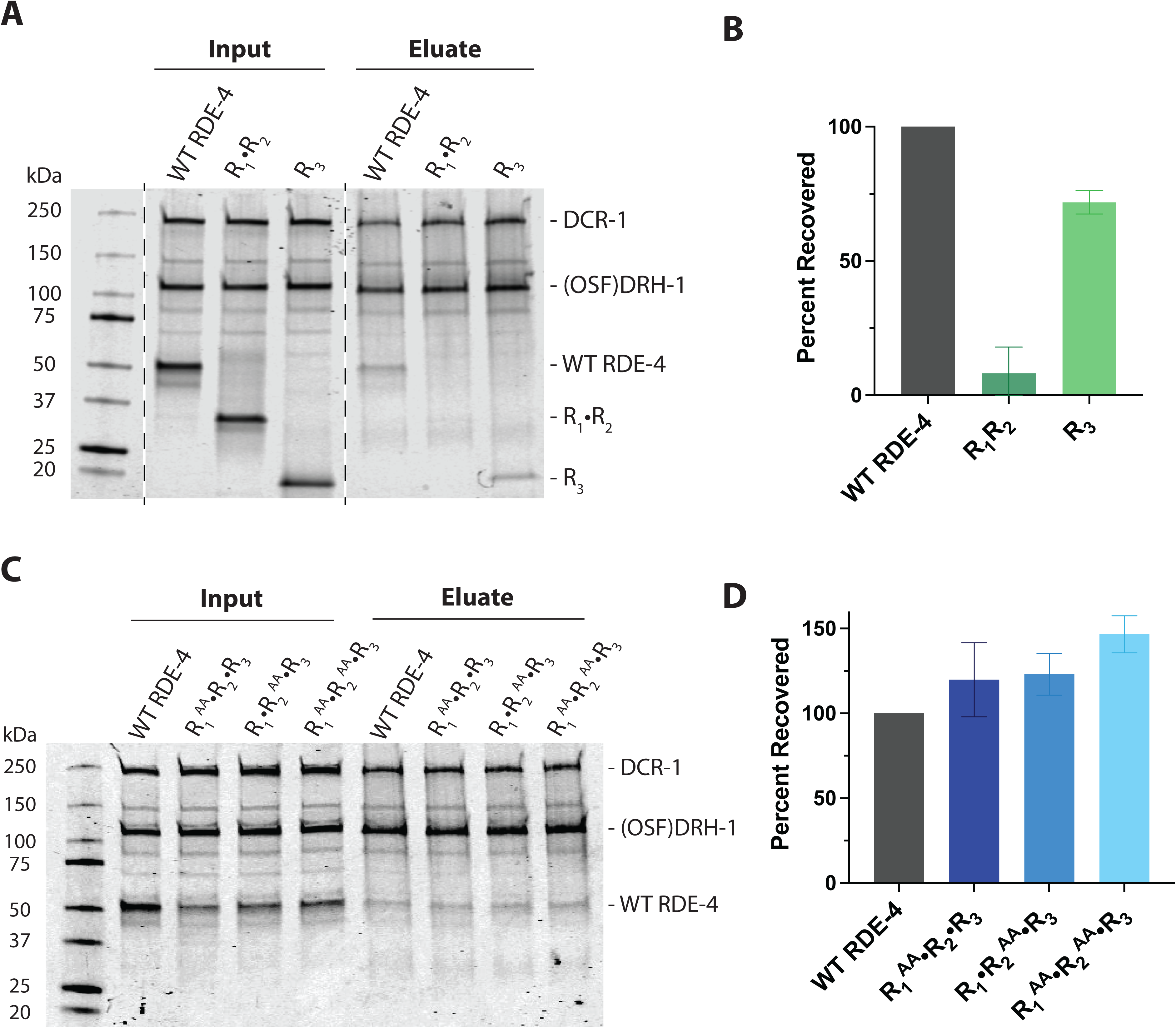
dsRBM3 is the primary driver for interaction with DCR-1•DRH-1. A. SDS-PAGE gel showing input (left, 9% of total sample) and eluate from StrepTactin beads (right, 100% of total sample) for DCR-1•(OSF)DRH-1 (146 nM) in the presence of WT RDE-4 or truncations (292 nM). Stained with SYPRO-red. Molecular mass markers (kDa), left side of gel; expected migration for proteins of interest, right side of gel. Gels were cut between lanes (as marked by dotted lines) to remove empty or irrelevant wells. No other image manipulations were performed. B. Data in panel A were quantified to determine amount of RDE-4 or truncations pulled down by DCR-1•(OSF)DRH-1. Values were plotted relative to amount of full-length WT RDE-4. Data points are mean ± SD (n=3). C. SDS-PAGE gel showing input (left, 9% of total) and eluate from StrepTactin beads (right, 100% of total) for DCR-1•(OSF)DRH-1 (146 nM) in the presence of WT RDE-4 or dsRBM point mutants (292 nM). Stained with SYPRO-red. Molecular mass markers (kDa), left side of gel; expected migration for proteins of interest, right side of gel. D. Data in panel C were quantified to determine the amount of RDE-4 or point mutants pulled down by DCR-1•(OSF)DRH-1. Values were plotted relative to the amount of full-length WT RDE-4. Data points are mean ± SD (n=3).

Additionally, point mutations in dsRBM1 and dsRBM2 showed >100% recovery, consistent with our hypothesis that RDE-4 affinity for DCR-1•DRH-1 is primarily through direct protein-protein contacts via dsRBM3 rather than dsRNA interactions (Figs. 5C-D). Importantly, pulldowns with dsRBM point mutants indicate that the reduced ATP hydrolysis and cleavage activity observed with R_1_•R_2_ ^AA^•R_3_ and R_1_ ^AA^•R_2_ ^AA^•R_3_ cannot be attributed to loss of complex formation.

To further characterize protein complexes present in our reactions, we used mass photometry to measure the mass and relative abundance of all species present in solution (Fig. 6 and Supplemental Table S1) (Cole et al. 2017; Soltermann et al. 2020; Young et al. 2018). This technique contributes to the knowledge gained from pulldowns because it is done with protein concentrations much closer to cleavage and ATP hydrolysis conditions, and because it allows visualization in a native context as opposed to denaturing PAGE. First, we analyzed the reconstituted antiviral complex with equimolar amounts of DCR-1•DRH-1 and WT RDE-4 revealing an array of protein complexes (Fig. 6A). On the far left of the trace was a peak attributed to free RDE-4, which has a theoretical mass of 46 kDa (Supplemental Table S1). The apparent mass was larger, likely reflecting an equilibrium between monomeric and dimeric RDE-4. The next peak to the right was attributed to free DRH-1 followed by a broad peak attributed to DCR-1•RDE-4. This broad peak is indicative of species heterogeneity likely because DCR-1 alone is unstable leading to a cluster of incomplete or partially degraded complexes. Notably, no strong signal was ever observed for a species the size of DRH-1•RDE-4. Finally, the peak on the far right of the trace represents DCR-1•DRH-1•RDE-4 and was used as a readout for antiviral complex formation. While the apparent mass was slightly smaller than the predicted mass, this is likely due to the presence of a DCR-1•DRH-1 complex that was included in the same peak, due to resolution limits of the instrument under these conditions, which can best distinguish differences of 50 kDa or more (RDE-4, ∼46 kDa). Once a mass distribution pattern was established with WT RDE-4, mutant RDE-4 constructs were added to DCR-1•DRH-1 under the same conditions (Figs 6B-F). This revealed that point mutations in dsRBMs 1 and 2 did not disrupt complex formation, as R_1_ ^AA^•R_2_ •R_3_ , R_1_ •R_2_ ^AA^•R_3_ and R_1_ ^AA^•R_2_ ^AA^•R_3_ all recapitulated the mass distribution of WT RDE-4 (Figs. 6B-D). Next, we tested our truncation constructs and found that the addition of R_3_ resulted in complete complex formation. In contrast, when R_1_•R_2_ was added to DCR-1•DRH-1 the largest visible mass was smaller than DCR-1•DRH-1 complex alone. This agrees with the idea that R_3_ promotes interaction between DCR-1 and DRH-1, a complex shown to be unstable without RDE-4 (Fig. 1D). However, it is worth noting that when the recommended parameters for protein concentration were exceeded in mass photometry, some species the size of the full antiviral complex could be seen upon addition of R_1_•R_2_. This observation agrees with our pulldown data, where higher input concentrations revealed residual interaction with the DCR-1•DRH-1 complex.

**Figure 6.**
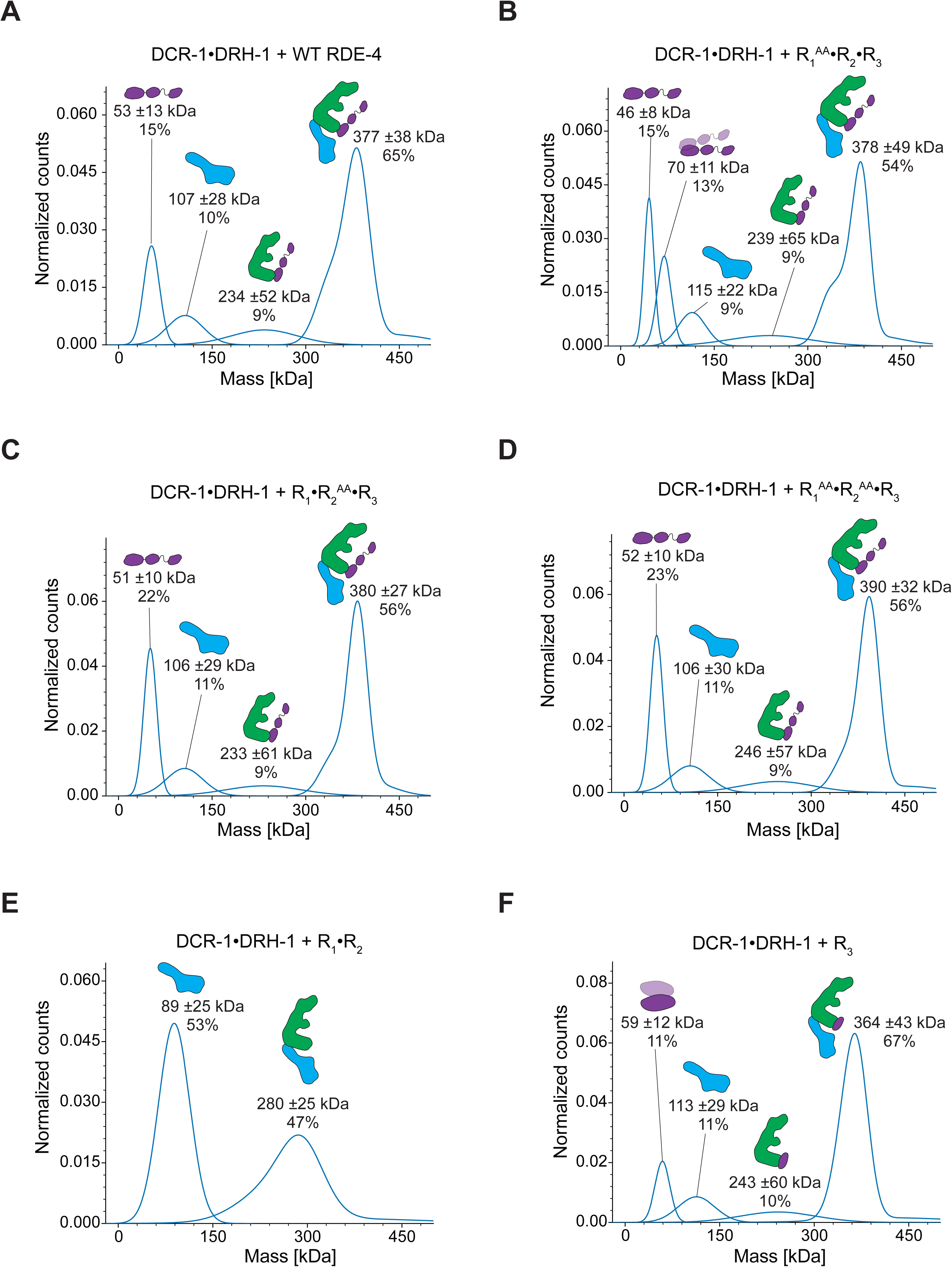
dsRBM3 promotes antiviral complex assembly. Panels A-F are representative Gaussian fits of mass photometry data (n=3). 100 nM DCR-1•DRH-1 and WT or mutant RDE-4 were combined. Various dilutions were tested using MP to optimize for counts and only those producing 500-5000 counts were retained. Peaks are labeled with apparent mass and standard error along with relative % abundance. Cartoons show prediction of protein complex that dominates labeled peaks.

### RDE-4 does not directly interact with DRH-1 alone

While mass photometry did not reveal a complex matching the mass of RDE-4 and full-length DRH-1, we sought further validation as this technique is best suited for relatively homogenous populations with mass differences of 50 kDa or more. Because DCR-1 and DRH-1 share a highly conserved helicase domain (Fig. 1A) where dsRBPs are reported to bind Dicer proteins, interaction between RDE-4 and DRH-1 could not be easily discounted. While DCR-1 cannot be purified alone to assay with RDE-4, DRH-1 can be tested more directly. Purification of full-length DRH-1 largely yields aggregated, inactive protein providing only a small amount of usable enzyme; however, truncation of the N-terminal domain (ΔNTD) produces a soluble, catalytically active form (Consalvo et al. 2024). We therefore assayed for interaction between RDE-4 and DRH-1 ΔNTD using pulldown experiments (Supplemental Fig. S4A). Equimolar WT RDE-4 and (OSF)DRH-1 ΔNTD were incubated in the absence of dsRNA, but no RDE-4 was detected in the eluate, indicating no stable interaction. Consistently, addition of RDE-4, even at an 8-fold molar excess, did not alter ATP hydrolysis by DRH-1 ΔNTD (Supplemental Figs. S4B-C).

## Discussion

Our reconstitution assays demonstrate that RDE-4 is not only sufficient to restore DCR-1•DRH-1 catalytic activity to levels observed with the co-purified antiviral complex DCR-1•DRH-1•RDE-4, but is also critical for maintaining complex stability. In the absence of RDE-4, DCR-1•DRH-1 rapidly loses activity, consistent with long-standing difficulties in purifying DCR-1 without binding partners (Consalvo et al. 2024). Addition of RDE-4 prevented this loss and allowed robust ATP hydrolysis and cleavage, suggesting that RDE-4 stabilizes both the integrity of the complex and its engagement with dsRNA. Importantly, pulldown and mass photometry experiments revealed that stable interactions occur primarily between RDE-4 and DCR-1, with no evidence of direct binding of RDE-4 to DRH-1 alone, and that these interactions are primarily mediated by dsRBM3 with some engagement by dsRBM2.

### Working Model

RDE-4 is abundant in cells and displays strong cooperativity, favoring coating of long, highly base-paired dsRNAs (Jannot et al. 2008; Parker et al. 2006, 2008; Thivierge et al. 2012). Thus, we propose a model in which RDE-4 binds viral dsRNA first (Fig. 7, Step 1). In this scenario, coating would not block DRH-1 binding, which recognizes termini, a property found in some other RLR family members (Consalvo et al. 2024). We speculate that translocation by DRH-1 in the DCR-1•DRH-1 complex is sufficient to displace bound proteins, generating local concentrations of unbound RDE-4 around DCR-1 and thereby promoting antiviral complex assembly (Aderounmu et al. 2025; Consalvo et al. 2024). This scenario is supported by observations that excess RDE-4 does not inhibit ATP hydrolysis or cleavage under the conditions tested in this study and suggests dynamic equilibria between dsRNA, RDE-4 and DRH-1 binding. However, it is also possible that RDE-4 remains bound to the dsRNA and directly recruits DCR-1•DRH-1 via dsRBM3. Indeed, AlphaFold predictions suggest RDE-4 primarily interacts with dsRNA via dsRBMs 1 and 2 leaving, dsRBM3 available for interaction (Supplemental Fig. S5) (Jumper et al. 2021; Varadi et al. 2024). In both models we predict that once the antiviral complex forms, RDE-4 is anchored to DCR-1•DRH-1 through the binding of dsRBM3 directly to DCR-1 (Fig. 7, Step 2); this is supported by pulldowns and mass photometry performed in this study, and previous cryo-EM data showing a single dsRBM of RDE-4 interacts with DCR-1’s helicase domain, although it was previously unclear which (Consalvo et al. 2024). We predict that interaction between DCR-1 and dsRBM3 of RDE-4 helps stabilize the interface between DCR-1 and DRH-1 so that when the full antiviral complex rebinds the stripped dsRNA, a large structural rearrangement occurs to generate a cleavage-competent conformation where both helicases simultaneously engage the same dsRNA. Once in this conformation, RDE-4 could aid in the positioning of dsRNA in the RNase III sites of DCR-1 through interaction with dsRBM2 (Fig. 7, Step 3). Similar mechanisms have been characterized for other RNase III enzymes where dsRNA actively engages the RNase III domains with assistance by dsRBMs (Gan et al. 2008; Nicholson 2014). By analogy, DCR-1 could act through a similar mechanism (Hansen et al. 2019). Once cleavage occurs, dsRBM1 may contribute by retaining siRNA products for handoff to RDE-1, ensuring efficient propagation of the silencing signal (Jannot et al. 2008; Knittel et al. 2025; Parker et al. 2008).

**Figure 7.**
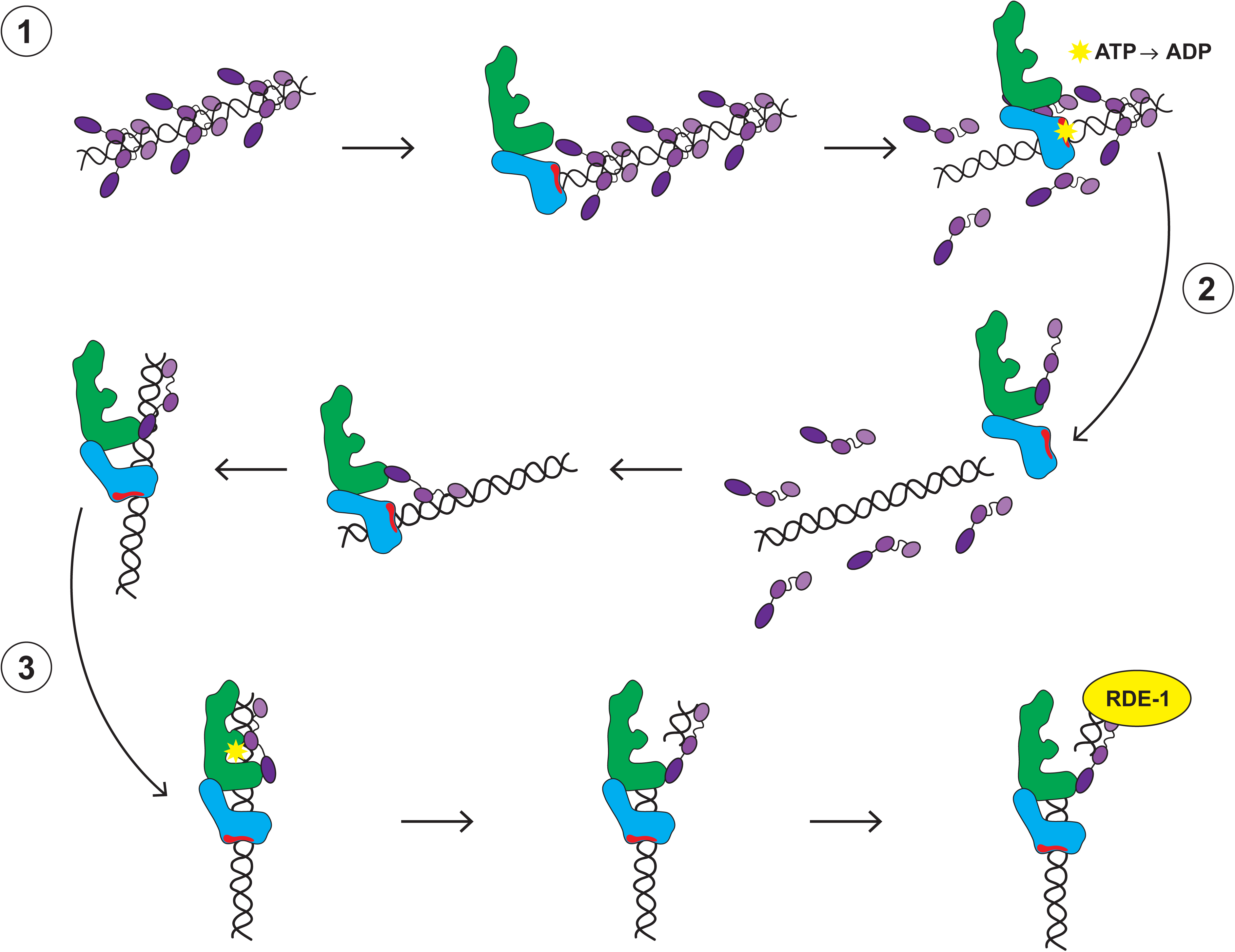
Working Model. Step 1 illustrates dsRNA substrate recruitment, step 2 illustrates antiviral complex assembly and step 3 illustrates cleavage and siRNA handoff. Steps are described further in the text. Purple indicates RDE-4 with different shades representing distinct dsRBMs. DRH-1 is shown in blue and DCR-1 in green with black dsRNA and yellow stars indicating ATP hydrolysis.

### Contributions of individual RDE-4 domains

Domain dissection revealed that dsRBM2 and dsRBM3 are essential for catalytic reconstitution, whereas dsRBM1 is dispensable. Our pulldown assays and mass photometry indicate dsRBM3 is primarily responsible for direct interaction with DCR-1 (Figs. 5A-B and Figs. 6E-F), anchoring RDE-4 to the complex. dsRBM2, by contrast, may play some role in mediating protein-protein interactions but is also the primary driver for dsRNA binding and potentially the positioning of dsRNA in the RNase III active sites. Point mutations within the KKxAK motif of dsRBM2 severely impaired both binding and catalysis, yet these constructs retained strong interaction with DCR-1•DRH-1 in pulldowns and mass photometry (Figs. 5C-D and Figs. 6B-D). Thus, we predict the primary role of dsRBM2 is not to stabilize protein-protein contacts but to orient dsRNA for productive cleavage. These findings refine earlier in vivo and extract-based models, where indirect stabilization by other cellular components could not be excluded (Blanchard et al. 2011; Parker et al. 2006, 2008). In contrast, our minimal system demonstrates that dsRBM2 and dsRBM3 act through distinct but complementary mechanisms: dsRNA positioning and complex anchoring, respectively.

The inhibition of cleavage by the R_1_•R_2_ truncation provides further mechanistic insight (Figs. 3C-D and Supplemental Fig. S3B). Despite retaining near-WT affinity for dsRNA (Fig. 4; Table 3), this construct strongly blocked cleavage, likely by coating dsRNA and occluding DCR-1 binding. We propose this coating cannot occur in the presence of the interaction between dsRBM3 and DCR-1. ATP hydrolysis was not inhibited by the R_1_•R_2_ truncation, likely because, in contrast to cleavage assays, ATP hydrolysis was monitored with excess dsRNA, and this served to titrate the inhibition by coating. This observation underscores the necessity of dsRBM3-mediated tethering; without anchoring to DCR-1, RDE-4 can bind dsRNA but cannot promote cleavage and instead acts competitively. We predict that this occurs because Step 2 in Figure 7 cannot take place, leaving the complex in a non-productive conformation where ATP hydrolysis can occur, but the complex cannot undergo the large structural rearrangement required to achieve robust cleavage. This relies on the assumption that both helicases can simultaneously bind the same dsRNA, and that this conformation is cleavage-competent (Consalvo et al. 2024).

### Length dependence of RDE-4-mediated activation

Our kinetic analysis revealed that RDE-4 activates both ATP hydrolysis and dsRNA cleavage by the DCR-1•DRH-1 complex, but the magnitude of this effect depends strongly on duplex length. For the 106-bp substrate, addition of WT RDE-4 increased ATP hydrolysis approximately 3-fold, with k_obs_ increasing from 0.048 to 0.13 min^-1^ and ν_0_ from 2.7 to 9.1 % hydrolysis/min (Table 2). Cleavage activity was similarly enhanced (∼2-fold), with k_obs_ increasing from 0.12 to 0.25 min^-1^ and ν_0_ from 9.4 to 16 % cleavage/min. It is worth noting that while both k_obs_ and ν_0_ are reported, some of these data, particularly conditions that failed to reconstitute or had low overall activity, like cleavage with 52-bp dsRNA, are not suitable for the exponential fit required to determine k_obs_, and are largely linear under these conditions. Therefore, ν_0_ is a more accurate reporter for trends observed across all constructs tested. Regardless, these results indicate that RDE-4 not only accelerated substrate turnover but also stabilized the complex to achieve a greater degree of catalysis, as indicated by the plateau (Table 1 and Figs. 3A-D).

On the shorter 52-bp substrate, RDE-4 still stimulated ATP hydrolysis, though the relative increases differed slightly (k_obs_ increased ∼2-fold and ν_0_ increased ∼4-fold), suggesting that DRH-1 activation by RDE-4 is largely length-independent. In contrast, cleavage showed a pronounced length effect: with WT RDE-4, ν_0_ for the 52-bp substrate was only 2.3 % hydrolysis/min compared to 16 % hydrolysis/min for the 106-bp substrate (Table 1; Figs. 3A-3D; Supplemental Figs. S2A-D). Thus, while DRH-1 hydrolysis responds robustly to RDE-4 on both duplexes, productive cleavage by DCR-1 is strongly length-dependent. We infer that longer substrates are required to accommodate simultaneous engagement of DRH-1 and DCR-1, consistent with our proposed cleavage-competent model (Fig. 1B).

This model is supported by recent biochemical and structural studies. For example, cryo-EM analysis of the DCR-1•DRH-1•RDE-4 complex bound to 52-bp dsRNA identified multiple conformational classes (Consalvo et al. 2024). A large fraction of particles showed high-resolution data for just the helicase domain of DRH-1, and while the rest of the complex was presumably present, it was blurry indicating a high level of conformational flexibility. Among the classes where resolution of DCR-1 was possible, dsRNA was frequently observed bound to DRH-1 but oriented away from the DCR-1 active sites, or bound at the DCR-1 PAZ/Platform domains in a configuration not aligned for cleavage. Notably, no classes showed a single dsRNA engaged simultaneously by both DRH-1 and the DCR-1. Together with our biochemical findings, these structural observations might suggest that while short dsRNAs can recruit DRH-1 and RDE-4, they are insufficient to promote the conformational state required for cooperative engagement of both helicase and RNase III domains, explaining the length dependence of cleavage efficiency.

### Conclusions

Our findings shed light on how *C. elegans* maintains antiviral activity using a single Dicer enzyme. Whereas *Drosophila* uses specialized Dicers (dmDcr-1 for miRNA and dmDcr-2 for siRNA), *C. elegans* relies on a single, multifunctional DCR-1 to process miRNA, endogenous siRNA and viral siRNA precursors, which are structurally quite diverse (Chen and Hur 2022). This versatility arises through collaborative interactions with accessory proteins including DRH-1 and RDE-4, in the case of antiviral RNAi. Throughout the evolution of Dicer’s helicase domain, there are notable decreases in dsRNA affinity and translocation activity (Aderounmu et al. 2025). However, by recruiting RDE-4, which has a high affinity for dsRNA, and DRH-1, a helicase protein capable of translocation, *C. elegans* has compensated for evolutionary changes to maintain a complex suitable for antiviral defense. This division of labor, anchoring and dsRNA recognition by RDE-4, ATP-driven scanning by DRH-1, and cleavage by DCR-1, provides a modular solution to the problem of substrate diversity (Aderounmu et al. 2025; Consalvo et al. 2024). Because humans, like *C. elegan*s, encode a single Dicer that is primarily dedicated to miRNA biogenesis (Wilson and Doudna 2013), elucidating how accessory proteins broaden the scope of DCR-1 substrates may offer clues to reprogramming human Dicer toward antiviral RNAi.

## Materials and Methods

### Plasmids and Cloning

*rde-4* pLib vectors were generated as described (Consalvo et al. 2024). Site directed mutagenesis was utilized to introduce point mutations or premature stop codons to generate truncated protein.

Sequences were confirmed using Plasmidsaurus. Mutations introduced in dsRBM1 were K89A and K90A, and for dsRBM2, K217A and K218A. Truncation for R_1_•R_2_ and R_3_ was at residue M244 (R_1_•R_2_ ends with D243 and M244 begins R_3_). Vectors were then transformed into DH10Bac *Escherichia coli* chemically competent cells to generate bacmid as described (Sinha and Bass 2017).

### Protein Expression and Purification

Purified bacmids were transformed into *Spodoptera frugiperda* (Sf9) insect cells and proteins expressed using the modified Bac-to-Bac Baculovirus Expression System as described (Sinha and Bass 2017) for both DCR-1•DRH-1 and RDE-4 constructs. Following large scale expression, DCR-1•DRH-1 was purified as described (Consalvo et al. 2024), except for the following modifications. Lysis buffer was supplemented with 0.5% Nonidet P40, instead of Triton X-100, and 200 µg/mL DNaseI Grade II. Following purification, the DCR-1•DRH-1 buffer (25 mM HEPES pH 7.5, 10 mM KOAc, 2 mM Mg(OAc)_2_, 100 mM KCl, 1 mM TCEP, 10% glycerol) was supplemented with an additional 20% glycerol for storage (30% glycerol total).

RDE-4 constructs were purified as described (Parker et al. 2006, 2008), with the following modifications. Lysis buffer was supplemented with 1% Triton X-100, 250 mg/ml DNase I Grade II, cOmplete EDTA-free protease inhibitor (1 tablet per 25 ml buffer), 0.7% (vol/vol) Protease Inhibitor Cocktail, and 1 mM phenylmethanesulfonyl fluoride (PMSF). Buffer A contained 30 mM HEPES pH 7.5, 1 mM TCEP, 10% glycerol, 30 mM imidazole and 500 mM NaCl, with successive washes containing 250 mM and 100 mM NaCl respectively. Buffer B contained 30 mM HEPES pH 7.5, 1 mM TCEP, 10% glycerol and 100 mM NaCl. All RDE-4 constructs, except R_3_, were found in the flowthrough of the heparin column and concentrated using a Vivaspin 20 concentrator. Size exclusion chromatography was then performed using the same buffer as DCR-1•DRH-1 (25 mM HEPES pH 7.5, 10 mM KOAc, 2 mM Mg(OAc)_2_, 100 mM KCl, 1 mM TCEP, 10% glycerol) and a HiLoad 16/600 Superdex 200 prep grade gel filtration column.

Concentration of DCR-1•DRH-1 and full length RDE-4 proteins following purification was determined using a NanodropOne and confirmed using SDS-PAGE and densitometry. The concentration of RDE-4 truncations R_1_•R_2_ and R_3_ were determined using SDS-PAGE and densitometry, due to the low number of aromatic residues (Supplemental Fig. S1C). Samples were run on SDS-PAGE alongside serial dilutions of BSA to use as a standard for densitometry. Gels were stained with SYPRO-Red, imaged with a Li-Cor Odyssey CLx, and quantified using ImageStudios software. Affinity purification tags were left on for recombinant proteins used in this study.

### dsRNA Sequences and Preparation

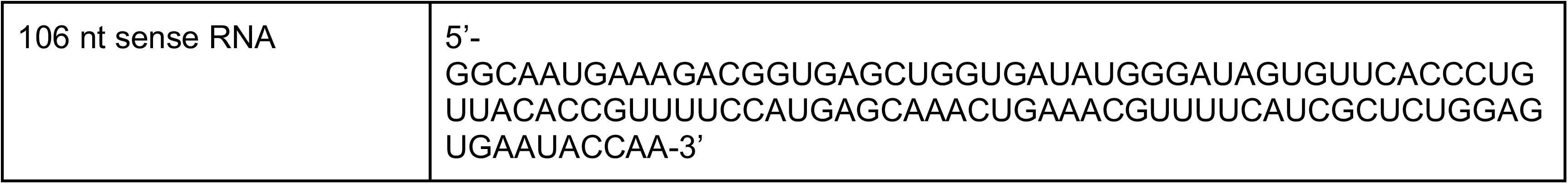
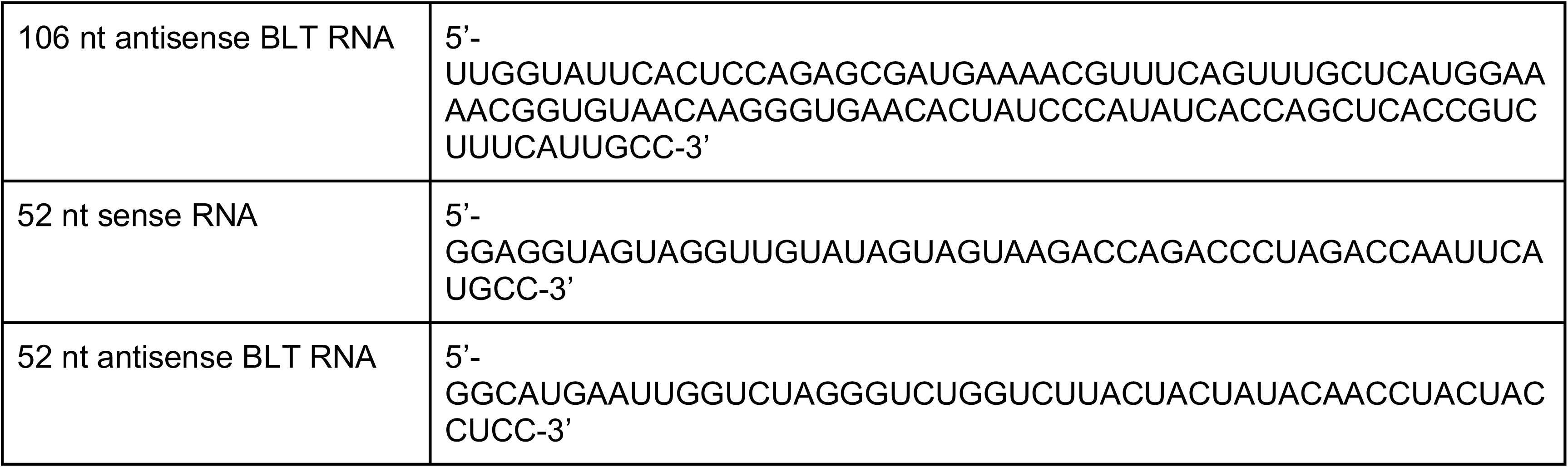

106-nucleotide (nt) RNAs were prepared as described (Consalvo et al. 2024; Sinha et al. 2015) except terminal cyclic phosphates were always removed following transcription. 52-nt RNAs were chemically synthesized by IDT and gel purified with denaturing PAGE (8% polyacrylamide (19:1), 8M urea, 1x TBE gel; 28W, 1.5 hrs). Equimolar amounts of ssRNAs were annealed in annealing buffer (50 mM TRIS pH 8.0, 20 mM KCl) by placing the reaction on a heat block (95°C) and slow cooling ≥2 hr. dsRNAs were then gel purified with 8% polyacrylamide native PAGE (8% polyacrylamide (19:1), 1x TBE gel; 8W, 1.5 hrs).

### ATP Hydrolysis Assay and Preincubation Assay

Assays were performed as described (Consalvo et al. 2024) with one modification: DCR-1•DRH-1 and RDE-4 were combined at the molar ratio listed and then preincubated at 20 °C for 1 minute before adding a master mix containing reaction buffer, ATP and dsRNA. When evaluating DCR-1•DRH-1 degradation, proteins were preincubated for 0-5 minutes before adding master mix. Exponential fits were attempted for all datasets; however conditions that failed to reconstitute WT activity were largely linear under these conditions making them less suitable for a single-exponential fit (used to determine k_obs_). For these cases, initial velocity (ν_0_) better represents relative activities among mutants and across substrates.

### Establishing Units of Activity

DCR-1•DRH-1 activity was quantified using 106-bp dsRNA. Because absolute activity levels varied between enzyme preparations, for each new purification, several dilutions were tested to determine the concentration that produced a plateau corresponding to 60% ATP hydrolysis and 80% dsRNA cleavage after 10 minutes. Absolute enzyme concentration was estimated from absorbance at 260 nm, yielding typical working concentrations of 100-150 nM. The activity obtained under these conditions was defined as 10 units of activity, representing the amount of enzyme required to achieve the specified plateau level of substrate conversion within standard assay conditions. The amount of RDE-4 added was adjusted to match units of activity so that a 1:2 molar ratio was maintained between DCR-1•DRH-1 and RDE-4. For example if 10 units of activity from one prep was equivalent to 150 nM DCR-1•DRH-1, then 300 nM RDE-4 would be added to the reaction; if 10 units of activity from another prep was equivalent to 100 nM DCR-1•DRH-1, then 200 nM RDE-4 would be added.

### dsRNA Cleavage Assay

Assays were as described (Consalvo et al. 2024) with one modification: DCR-1•DRH-1 and RDE-4 were combined at the molar ratio listed and then preincubated at 20 °C for 1 minute before adding a master mix containing reaction buffer, ATP and dsRNA. Exponential fits were attempted for all datasets; however conditions that failed to reconstitute WT activity were largely linear under these conditions making them less suitable for a single-exponential fit (used to determine k_obs_). This was also true for all complexes tested with 52-bp dsRNA, where the amount of cleavage was very low across WT and mutants. For these cases, initial velocity (ν_0_) better represents relative activities among mutants and across substrates.

### Pulldown Assay

2 µg of (OSF)DRH-1 (146 nM DCR-1•DRH-1) was combined with 2-fold molar excess of WT or mutant RDE-4 (292 nM). Proteins were incubated together at 4°C for 30 minutes before adding to pre-washed Cube Biotech PureCube HiCap StrepTactin MagBeads. 20 µL beads were washed 3x with binding buffer (25 mM HEPES pH 7.5, 10 mM KOAc, 2 mM Mg(OAc)_2_, 100 mM KCl, 1 mM TCEP, 10% glycerol) prior to addition of protein. Protein was then incubated with beads at 4°C for 2 hours (mixing) then washed 2x with binding buffer before adding 25 µL elution buffer containing 3.5 mM desthiobiotin. Beads and elution buffer were incubated at 4°C for 1 hour (mixing). 9% of input and 100% of elution were then analyzed using an SDS-PAGE gel stained with SYPRO-red. The fraction of RDE-4 recovered was normalized to amount of DRH-1 present in the elution. This protocol was repeated using 2 µg (OSF)DRH-1 ΔNTD (190 nM) with 2-fold molar excess WT RDE-4 (380 nM).

### Gel Shift Assay

0-51 µM WT or mutant RDE-4 protein was incubated with 20 pM ^32^P-end-labeled 106 BLT dsRNA for 30 min at 4°C. Binding was visualized with native gel electrophoresis. The gel contained 8% polyacrylamide (29:1), 1x TBE and was run in 1x TBE at 4°C. Gel was dried for 1 hour, exposed overnight, and imaged using a PhosphorImager. Calculations for dsRNA binding affinity (K_d_) used total protein concentration without mathematical correction for protein dimerization.

### Mass Photometry

DCR-1•DRH-1 and RDE-4 were combined in an equimolar ratio (100 nM). Droplet-free focusing was performed and 20 µL of sample were added to the Refyen TwoMP using microscope cover slides and silicon gaskets from Refeyn Ltd. Dilutions (1:2, 1:3 and 1:5) were also measured using the droplet-dilution focusing method to optimize for ∼2000-6000 total counts per sample. The sample chamber was prepared, and data were acquired as described (Wu and Piszczek 2021). Events were imported and histograms generated using DiscoverMP software. Histograms were auto-fit with Guassian curves (rather than manually selecting regions), and a representative trace for one replicate is shown. Peaks are labeled with mass assigned by instrument, and proteins predicted to correspond with these masses are depicted as cartoons. Theoretical protein masses are summarized in Supplemental Table 1.

## Acknowledgments

We thank the Bass lab for helpful feedback and the Shen lab for assistance with Mass Photometry, supported by the Utah Research Instrumentation Fund. DNA synthesis and flow cytometry was performed at the University of Utah Core Facilities. Proteomics mass spectrometry analysis was performed at the Mass Spectrometry and Proteomics Core Facility at the University of Utah. Mass spectrometry equipment was obtained through a Shared Instrumentation Grant #S10 OD018210 01A1. This work was supported by funding to B.L.B from the National Institute of General Medical Sciences (R35GM141262). B.L.B. is a Jon M. Huntsman Presidential Endowed Chair.

## Notes

### Competing Interest Statement

The authors have declared no competing interest.

